# Improving reproductive success in captive marmosets through active female choice

**DOI:** 10.1101/2024.05.08.593247

**Authors:** Taylor M Drazan, Sean P Bradley, Aikeen Jones, Krystal Allen-Worthington, Yogita Chudasama

## Abstract

The recent upsurge in the use of common marmosets (Callithrix jacchus) as a desirable model for high priority biomedical research has challenged local and global suppliers struggling to provide sufficient numbers of marmosets for large scale projects. Scientific research laboratories are increasingly establishing institutional breeding colonies, in part to combat the resulting shortage and high cost of commercially available animals, and in part to have maximum control over research lines involving reproduction and development. For such laboratories, efficient marmoset breeding can be challenging and time consuming. Random male/female pairings are often unsuccessful, with intervals of several months before attempting alternate pairings. Here we address this challenge through a behavioral task that promotes self-directed female selection of potential mates to increase the efficiency of breeding in captive marmosets. We created a partner preference test (‘love maze’) in which nulliparous females (n=12) had the opportunity to select between two eligible males (n=23) at a time, in a forced choice test. In this test, both males usually displayed sexual solicitations. However, the female would clearly indicate her preference for one. Most commonly, the female actively ignored the non-preferred male and directed overt prosocial behaviors (e.g. proceptive tongue-flicking, approach and grooming) to the preferred male. Moreover, once a male was selected in this context, the female would continue to prefer him over other males in three consecutive testing sessions. Compared with random pairings, this directed female choice showed a 2.5-fold improvement in breeding within 90 days compared to random pairings. This cost-effective and straightforward pairing practice can be used to enhance breeding efficiency in both small and large marmoset colonies.

## Introduction

The common marmoset (Callithrix jacchus) has figured prominently in biomedical research (1–5), and recently ‘re-discovered’ as a model organism for its unique reproductive and developmental traits (6,7). In particular, young infant marmosets serve as very useful models for understanding putative developmental abnormalities that may impact the normal expression of social, cognitive, and emotional behavior. Similar to humans, marmosets live in close-knit families with a high degree of cooperative care and learn through social and vocal interactions (8,9). In addition, critical features of the primate brain, such as the prefrontal cortex, temporal lobe and thalamus, and their connections, are shared between the marmoset and the human (10,11). Thus, this small primate can model important changes in disorders of human affective and cognitive state, and accordingly, is being used on an increasing scale for anatomical, physiological, genetic and behavioral research.

The enthusiasm for marmosets as a nonhuman primate model for biomedical research has challenged suppliers across the globe unable to provide sufficient numbers of marmosets for large scale projects (12,13). Coupled with the high cost of purchasing marmosets from commercial vendors which currently stands at approximately $12,000 USD per monkey, many laboratories have resorted to creating their own breeding colonies. However, while marmosets are hardy and live for many years in captivity, breeding them in the laboratory presents certain difficulties (14). For example, although marmosets breed cooperatively, most fully mature subordinate females fail to reproduce due to ovulatory suppression, while others attempt to breed but produce very few surviving infants (15–17). Sometimes, directed aggression or harassment from a dominant or rival female can interfere with breeding attempts by inhibiting ovulation, and can induce spontaneous abortion, as has been observed in other species (18–20).

In captivity, male/female marmoset pairings for breeding are usually determined based on age, health measures, and history of reproductive success. These factors are balanced with a need to maintain an outbred population with the aim of creating as much genetic variation as possible. The creation of new male-female breeding pairs is determined gradually over several weeks of introductions to ensure partner compatibility. However, even taking these factors into account, the time from mating to conception can range from 140 – 530 days (7,14) resulting in a breeding rate that is less than optimal. If paired couples prove incompatible due to fighting or aggression, the female is introduced to another male and several pairings might ensue before a compatible match is found. Arranged partnerships of this kind can be stressful for the animal and are also labor, intensive requiring many hours and days of careful observation, and even if there is compatibility, there is no guarantee of reproductive success. Moreover, although the male can influence the females’ reproductive success by offering her resources and benefits such as food and rank, in captivity, some females fail to conceive despite successful co-habitation with high-ranking males. In some cases, otherwise cooperative marmoset pairs live together for months or years but fail to conceive.

Here we address the possibility of increasing the efficiency of breeding within a marmoset colony by introducing a systematic means of self-directed female choice of potential mates. We exposed potential pairs through a novel partner preference test (known colloquially as the “love maze”), in which behavioral cues indicated their preference to mate, in a forced choice social paradigm. With multiple runs of this test, each female had the opportunity to select between 3-6 eligible males, and we proceeded with pairings based on this selection. We found that this practice significantly improved the success of pair mating’s and decreased the latency to conceive.

## Results

### Marmosets display sociosexual behaviors in partner preference love maze

Our test subjects were 12 nulliparous females (age 2–6yrs). Twenty-three males (age 2-7yrs) were required to test the female’s preference for mates as breeding partners. We first created a partner preference test (known colloquially as the ‘love maze’) using custom designed apparatus to allow female marmosets to choose between two males, and exhibit behaviors that could be viewed as preferentially directed toward one male over the other (**Fig. 1A).** Along with other callitrichine primates, marmosets display selective social and sexual preferences towards a specific mate manifesting behaviorally as a pair-bond (21). Interactions with potential sexual partners involve displays of discrete affiliative and sexual behaviors which include tongue flicking, body presentations, lip-smacking, and allogrooming (22,23).

**Figure 1.**
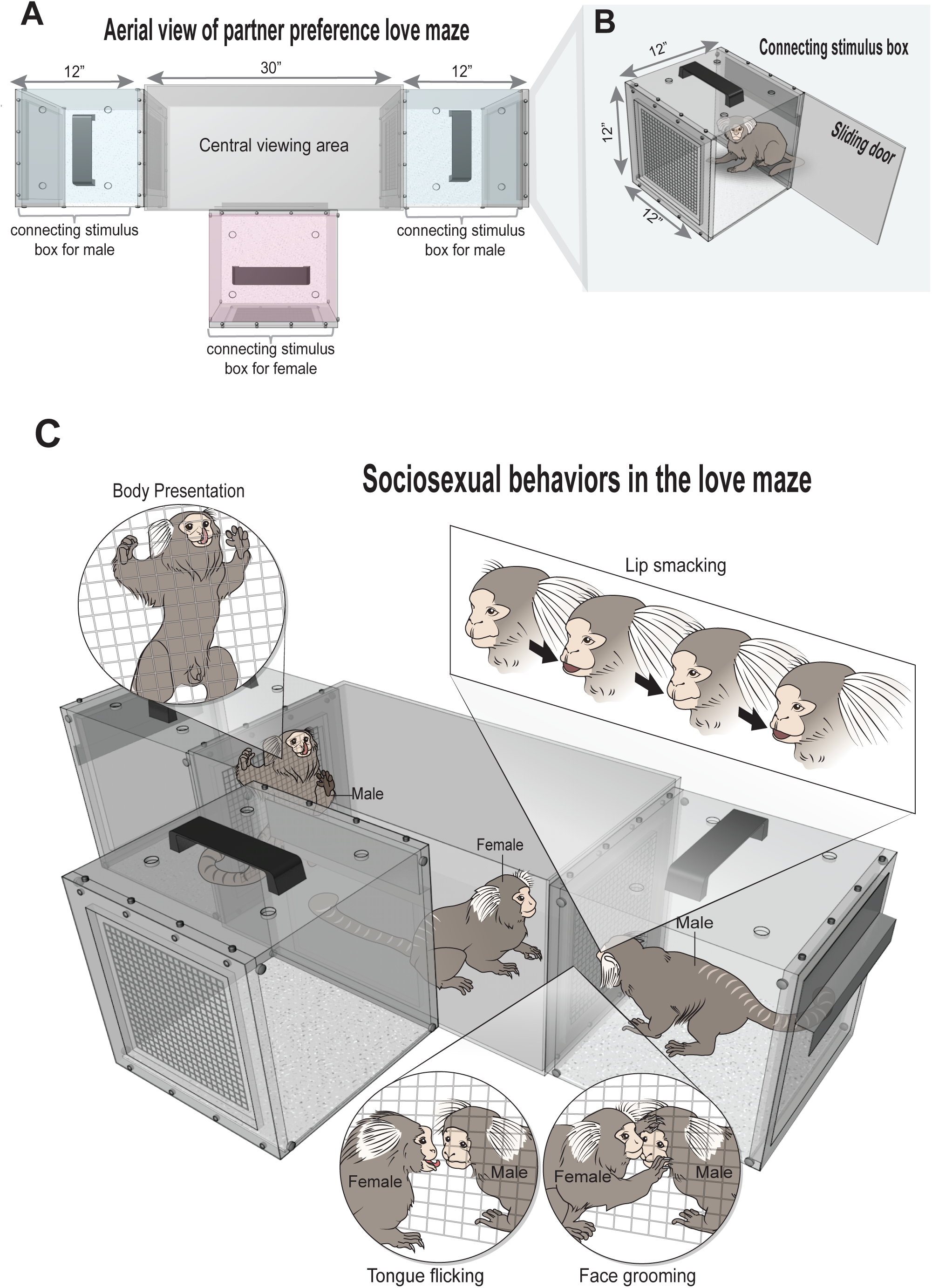
Partner preference love maze apparatus and sociosexual behaviors. (A) Aerial view of apparatus showing T-maze configuration. (B) Schematic of connecting stimulus box with sliding door to allow monkey to enter and exit the box. The stimulus box connects with the home cage and central viewing area of the partner preference apparatus. (C) Schematic illustration of sociosexual behaviors. Both males and females displayed sexual solicitations. The central viewing area provided a safe zone for the female to sample, observe and interact with the two males without being subject to injury or aggression. The preferred male was solicited with tongue flicks, face grooming and lip smacking. The female also sat in close proximity to her chosen male. Sometimes, the non-preferred male vied for her attention with body presentations or vocalizations even though her attention was focused on the other male.

To observe sexual solicitations, the partner preference love maze was made of Plexiglass and configured like a T-shaped maze (**Fig. 1A**). Animals voluntarily entered a custom designed stimulus box adapted to mount securely onto the home cage **(Fig 1B**). A sliding door allowed the animal to enter and exit the box without the need for experimenter handling. The stimulus box containing the marmoset was then safely transported to the testing apparatus and securely attached to the love maze apparatus. The female was placed in the center and two males, both unknown to the female, were placed in the right and left flanking positions (**Fig 1C**). Only the female was allowed to freely explore the central viewing area and approach either male depending on her preference. The males could see, hear, smell, and lightly touch the female through a wire mesh only. If the female was genuinely interested in one male over the other, she was required to confirm this preference by displaying sexual solicitations towards that male on another two separate sessions during which her preferred male was presented alongside a different unknown male (**Fig. 2A**).

**Figure 2.**
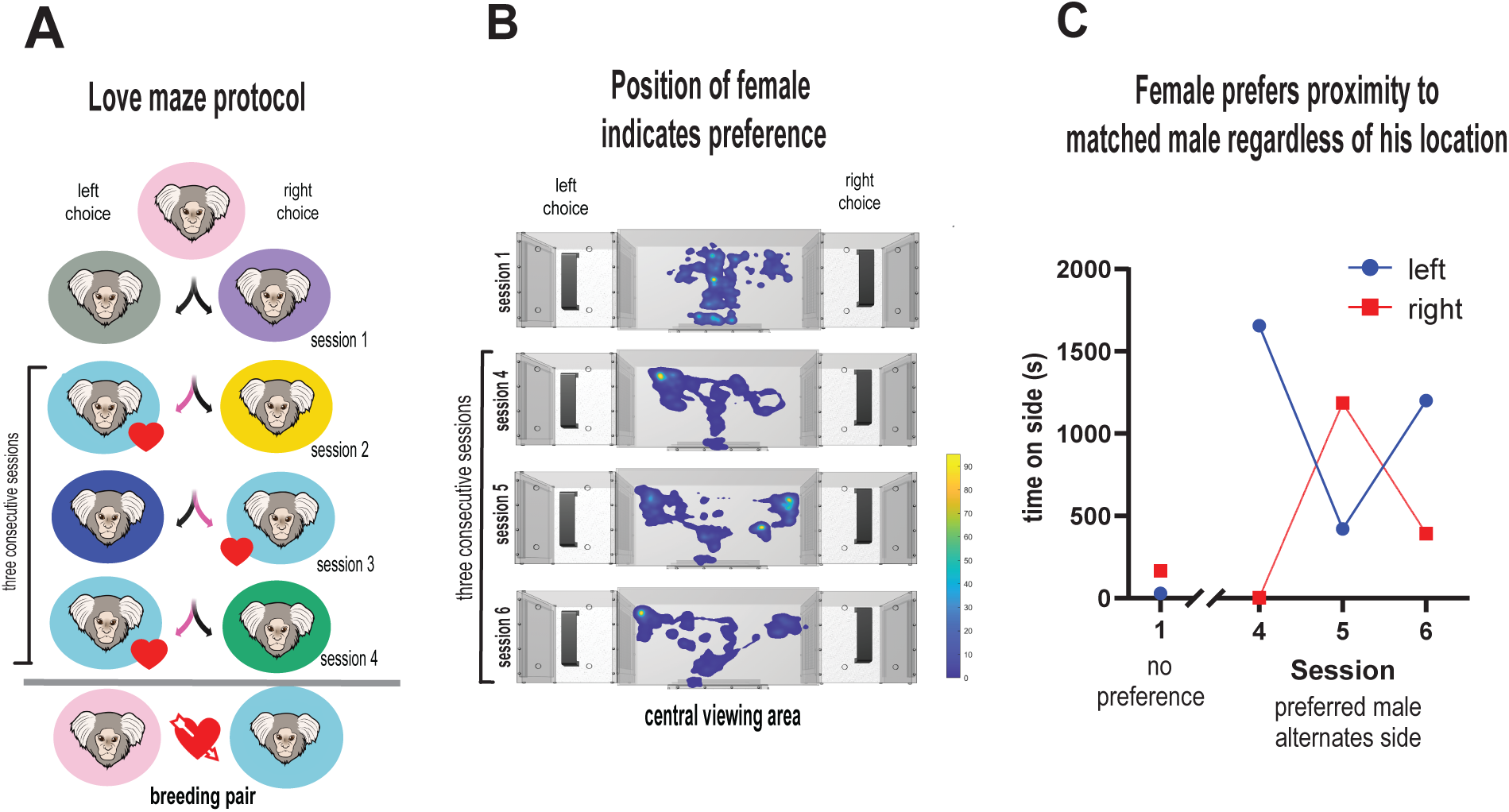
Love maze protocol. (A) In the love maze protocol, the female is given the opportunity to sample two unfamiliar males by exhibiting sexual solicitation toward one male. A schematic marmoset monkey face superimposed on a pink circle indicates the choosing female. All other colors represent different males that were offered as pairs. The chosen male is depicted by a monkey face superimposed on the light blue circle, a red heart and pink arrows directed towards the preferred male. (B) Representative heat maps, superimposed on central viewing area, of a single female indicated her preference . Hot colors on the heat map indicate greater preference due to close proximity of the female to that male in that location. In her first session (top panel), the female sampled both males but did not show any preference. On sessions 2 and 3, she showed a preference to one male (data not shown), but when that male was presented alongside another male on session 4, she shifted her preference to the ‘new’ male. She then continued her preference for the new male on sessions 5 and 6. Three consecutive sessions of choosing the same male were considered a match for a breeding pair. (C) Time spent in close proximity to the preferred male by female represented in panel B. The left/right location of the preferred male was alternated between sessions. Consistent with the heat maps, she identified her preferred male and followed his left/right location.

In addition to displaying sexual solicitation behaviors, the female’s decision to stay in close proximity to the male was another indicator of her mate of choice. We tracked her movements in the central viewing area and visualized her activity data as heat maps (**Fig. 2B**). The heat maps revealed that the female sampled both males by approaching each male in their respective locations. When she was not interested in either male, most of her activity was restricted to the central zone. Over the sessions, she developed a preference for one male by positioning herself in close proximity to him. Moreover, because we alternated the left/right location of the preferred male between sessions, she readily identified her preferred male in his new location and stayed in close proximity to him in that new location (**Fig. 2C).** Thus, once her decision was made, most of her time was spent in close social contact with her preferred male.

Altogether, the love maze apparatus and protocol provided the female an opportunity to sample her choice of males in a secure and safe environment without the risk of aggression, while directing her preference for a single partner on repeated occasions. Of our original sample of 12 nulliparous females, two females were highly aggressive towards all males and could not be paired leaving a female sample size of n=10. Three sessions of new male introductions were usually sufficient for the female to make her preferred choice. Five females required an additional 2 or 3 sessions with ‘new’ male options before she finalized her preference over three consecutive sessions (**see Fig. 2A,B**).

### Females exhibit sociosexual preferences for specific males

We observed a range of sexual solicitations exhibited by both males and females during the love maze protocol. Two independent raters observed the frequency and duration of behaviors during two separate sessions with an inter-rater reliability of >0.94 (p <0.01). On seeing the female, both males displayed obvious sexual solicitations including full body presentations with erection, as well as proceptive tongue flicks and lip-smacking behavior expressing affiliation towards the female **(Fig 1C**; **Table 1**). Both males and females exhibited almost equal amounts of sexual solicitations (t_(25.334)_ =-0.44, p > 0.05; **Fig 3A)**, but the number of sociosexual behaviors displayed by each animal varied substantially across age. Older animals of 5-6 years of age displayed more sexual behaviors in general (>50), but many 2-3 year olds were also expressive in their sexual displays even though they exhibited fewer behaviors (>30). In many cases, the expressed sexual behavior in younger animals was equivalent to the average 5-year-old (**Fig. 3B**). Moreover, apart from one older female who chose a young male, we found that almost all females chose older males, with a mean difference of -1.6 years (t(_9_) = -2.30, p < 0.05), presumably because their behaviors indicated to the female that those males were sexually experienced (**Fig. 3C).** However, despite being older, the preferred male was also lower in weight suggesting that they were smaller in size relative to the choosing female. On average, females tended to select male partners who were 70.2 g lighter than themselves (t(_9_) =2.68, p < 0.05; **Fig. 3D).**

**Figure 3.**
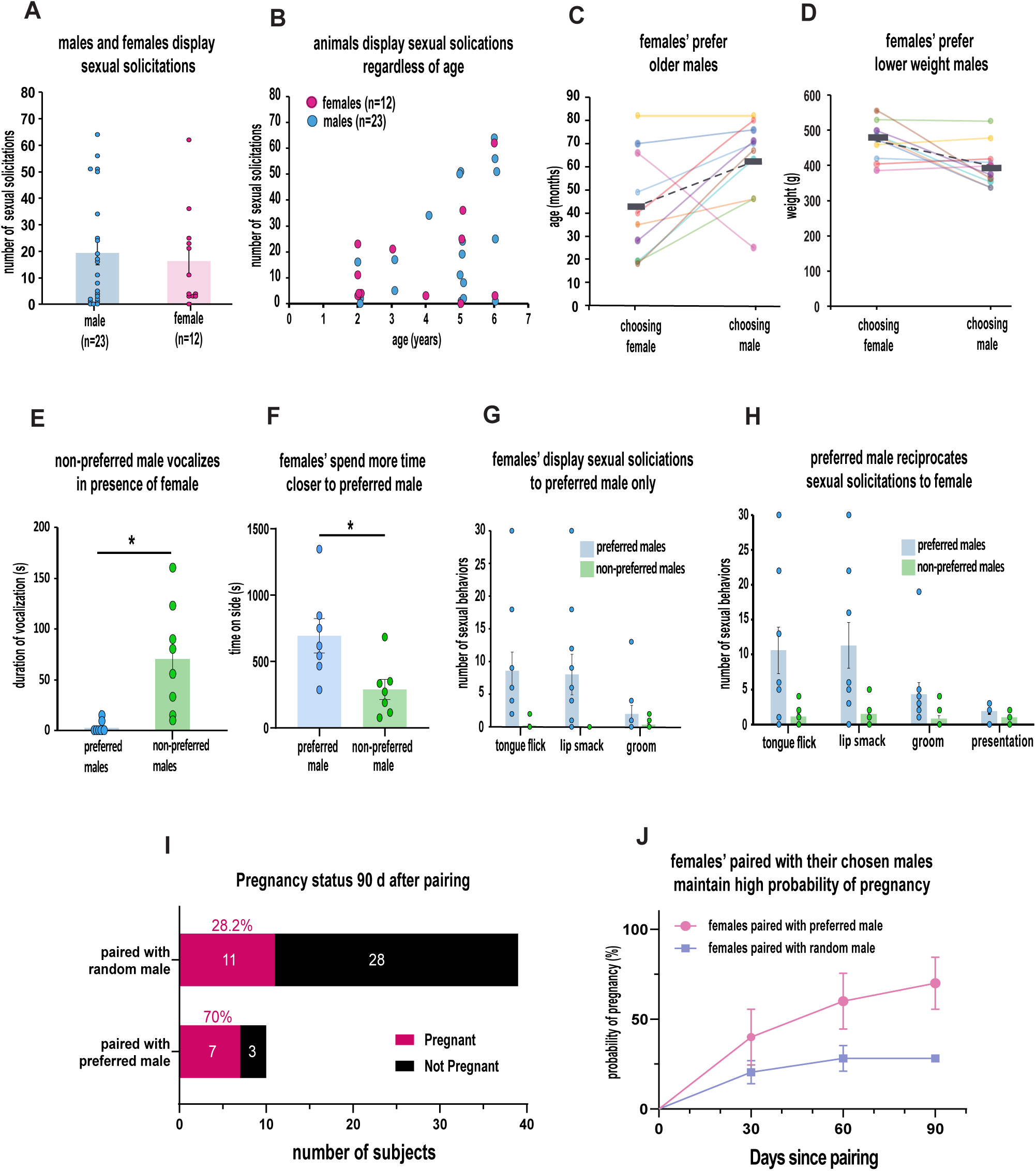
Females exhibit sociosexual preferences for specific males. (A) Males and females exhibit almost equal number of sexual behaviors on average. Circles represent individual animals (B) Scatterplot showing average number of sexual solicitations exhibited by all males and females as a function of age. (C) Line graph to show that female typically chose older males. Age of female is matched to the age of her chosen male. Each pair is represented by a different color. The gray dashed line represents the mean. (D) Line graphs to show that chosen male was lower in weight than the female. The weight of the female is matched to the weight of her chosen male Each pair is represented by a different color. The gray dashed line represents the mean. (E) The average duration of vocalizations during one session emitted by the non-preferred male while vying for the females attention. Circles represent scores of individual animals. (F) Females’ spent more time in close proximity to their preferred male. Circles represent scores of individual animals. (G) Females exhibit behaviors exclusively for the preferred male. Number of each sexual behavior exhibited by female toward the preferred and non-preferred males across three consecutive sessions. Circles represent scores of individual animals. (H) The preferred male reciprocates sexual behaviors to the female. Number of each sexual behavior exhibited by preferred and non-preferred males towards female. Circles represent individual animals. (I) Percentage of females that conceived 90 days after pairing with their chosen male relative to random male. (J) Probability of pregnancy in females paired with chosen male or random male as a function of days since pairing. Females paired with their chosen male conceive more and faster such that more marmosets are pregnant by 90 days.

**Table 1.** Description of sociosexual behaviors exhibited by both male and female marmosets

| **Observed behavior** | **Description of each behavior** |
| --- | --- |
| Tongue flicking | A rhythmic tongue-protrusion display while mouth is open. |
| Lip smacking | Mouth is pursed open and shut, usually very rapidly. |
| Grooming | A social interaction in which one monkey sits in close proximity to another and engages in touching or stroking the other monkey. Grooming of the body can last from seconds to several minutes. |
| Presentation | The monkey stretches its entire body to present its chest, stomach, or genitals towards another animal. The presenting animal might occupy different positions such as lying on the back or standing vertically while exposing its torso. |

Since the two males were facing each other, they became highly competitive in drawing the female’s attention. When the female directed her attention towards her preferred male, the non-preferred male expressed a staccato of aggressive or angry ‘chatter’ vocalizations (**Fig. 3E**) presumably to grab her attention towards him. The female, however, was very clear in her choice of male. In addition to spending more time in close proximity to her preferred male to encourage pair bonding between them (t(_6_) = 2.87, p < 0.05; **Fig. 3F**), she expressed more sexually directed behaviors towards him on multiple occasions. Proceptive tongue-flicking flicks (t_(9.1)_ =2.95, p < 0.05), a well-known invitational behavior in this species, and lip-smacking (t(_9_) =2.59, p < 0.05), a submissive signal that signifies affiliation, were the primary indicators of her chosen male (**Fig 3G**). Although grooming behavior was observed on multiple occasions, it was not a primary indicator of her preference (t_(9.5)_ =1.31, p = 0.05). The preferred male reciprocated sexual solicitations to the female thereby confirming her advances mostly with tongue flicks (t_(9.3)_ =2.81, p < 0.05) and lip smacks (t_(9.4)_ =2.96, p < 0.05), but less so with grooming (t_(10.2)_ =1.99, p > 0.05) and presentation behavior (t_(15.9)_ =1.71, p > 0.05; **Fig. 3H**).

### Female choice of male determines her reproductive success

The partner preference love maze allowed the female to choose her preferred mate in a safe environment. Our data show that in captivity, this choice is a major determinant of her reproductive success. We calculated the number of days it took the females to conceive with their preferred males relative to those females that were randomly paired with unknown males. Unfortunately, two of the twelve nulliparous females when paired with their chosen mates failed to conceive due to inherent fertility issues (i.e., polycystic ovaries). These two animals were removed from subsequent statistical analyses. The remaining ten females that were paired with their preferred male showed a 70% success rate in pregnancy within 90 days relative to only 27.5% in those male-female couples (n=40) that were randomly paired for breeding, representing a 2.5 fold increase in reproductive success within that interval (χ²(1) = 5.982, p <0.05; **Fig 3I**). Our data suggest that, for random male-female marmoset pairings, the success of conception did not improve much after the first month but the ten females matched with their preferred male exhibited a positive trajectory of reproductive success (χ²(1)= 4.818, p <0.05; **Fig. 3J)**. Four females that were paired with their chosen males in the love maze successfully conceived within 30 days. Since frequencies of proceptivity correlate positively with hormonal levels of testosterone and oestrone during the reproductive cycle, it is possible that the females that conceived quickly were indeed hormonally primed (24,25). The possibility remains, however, that had she been paired with her non-preferred male, despite his intense sexual solicitations towards her, she might not have conceived, taken a substantially long time to conceive or need to be separated due to aggression.

## Discussion

In this study we introduce a safe, and time efficient partner preference protocol using a low-cost and versatile piece of apparatus for establishing successful breeding pairs with a high rate of pregnancy. We found that despite the artificial setting of the partner preference love maze, both males and females readily engaged in sociosexual behavioral cues as a preference to mate, and females used these cues together with social contact to direct her mating choice. Importantly, the central viewing area offered the female safe exploration to sample both males at her leisure, without threat of aggression or injury which might occur when introducing two animals unfamiliar with each other. Given the opportunity to choose her breeding partner, we found these female marmosets had an advantage of conceiving as early as 30 days after pairing. We believe our cheap and easy practice will foster the growth of self-sustaining marmoset colonies by significantly lowering the time invested in finding compatible breeding pairs.

In captive breeding colonies, marmosets form long-term socially monogamous relationships based on mated pairs that are successfully caged together (26,27). In natural settings, marmosets are known to be flexible in their sexual preferences showing a willingness to mate with a potential mate in the absence of their pair-mate (28,29). This is especially true for the alpha female who retains reproductive sovereignty over subordinate females, and for whom promiscuous mating serves advantageously to reproductive success (26,30). Moreover, in most primate species, females do not solicit males, and when they do, they do not prefer only one male (31,32). Our data suggest that this pattern differs in socially housed captive female marmosets, including subordinates whose breeding is restricted due to ovulatory suppression (33). In the partner preference test, we found that females selectively chose one male whilst ignoring others, despite their active solicitations towards her. Notably, her preference correlated with her reproductive success. In primates, the usual metrics of female reproductive success, such as number of mounts or copulations, are not good indicators of pregnancy (34). We defined reproductive success as conception, detected through ultrasonography. This measure quantified the relationship between choice of male and mating success. In marmosets, the selective basis for female attractiveness relies on her reproductive experience and her rank, both of which make her attractive to males, thereby determining her reproductive success. Our data suggest that the strong solicitation by the female of an unknown male, highlights the importance of female choice in her reproductive success.

The precise social and sexual factors that might have contributed to the females’ selection is unknown. Marmosets exhibit little or no sexual dimorphism in body size which make it hard to identify age or rank. Moreover, the females in the present study had no direct interaction with males outside their own family group and had never been paired or interacted with the males they were offered. There was also a general tendency for the females to choose older males presumably because age accrues a selective advantage. However, there is a dearth of data on what constitutes attractiveness in captive marmosets. Early studies confirmed that age of male does not correlate with reproductive success (35,36), but since males are the principle caretakers of the young, it has been suggested that older males may be attractive because they are more experienced with cooperative care and invest more time with offspring (37). Importantly, although females chose males older than themselves, the older males were not ‘larger’ in size, an attribute of fitness that might indicate high rank or experience. We found that females typically chose males that were lower in weight than themselves, and one large 5-year-old female (499g) chose a young, and smaller 2-year-old (374g) male. If weight or size is construed as an indicator of rank, then the females in this study typically chose low-ranking males. This is not to say that these males would not be high-ranking in a different social context where they might compete for the same female. Abbot and Hearn (1978) reported that mated pairs comprising older females and young males provided the best combination of raising offspring. Low-ranking males provide more affiliative, social support of the offspring which can be incompatible with high-ranking males engaged in high competition (26). Our data are consistent with this view.

It is feasible that the safety imposed by the partner preference test provided unhindered freedom and motivation for the female to choose the less dominant or lower-ranking male. This choice might not be an option in natural settings where visible behaviors such as aggressive interactions take stage. While some labs have successfully paired two males with a single female to facilitate breeding, the female still favors one male over the other (33,38,39). Thus, if anything, random pairings of females with smaller or lower weight males in captivity are likely to yield better mating success. We recognize, that despite pairing the female with a partner of her choice, there is no guarantee that her offspring will survive, especially to first time mothers where pregnancy loss is common (14). However, if older females are experienced with social rearing in family settings, and younger males are socially affiliative, then successful breeding with high infant production is ostensibly possible.

Marmosets are being used on an increasing scale for biomedical research partly because of their reproductive biology and compatibility with gene editing techniques (6). As such, the marmoset provides a strategic choice for longitudinal developmental and transgenic studies which require thriving breeding colonies. While most studies focus on the qualities of successful males, as the stronger selective force in determining reproductive success, we show that dominant features of rank and size may not always feature in her decision, and that her choice of smaller, less dominant monkeys may be the critical requirement for a successful marmoset mating system. The partner preference ‘love maze,’ provides a time efficient means of achieving this goal.

## Materials and Methods

### Subjects

A total of 35 adult common marmosets (Callithrix jacchus), ranging in weight from 250-600g at the start of behavioral testing were used in this study. Twelve nulliparous females (age 2–6yrs) served as the test subjects. Twenty-three males (age 2-7yrs) were required to test the female’s preference for mates as breeding partners. They were housed in temperature - controlled rooms (76–80°F) under diurnal conditions (12-h light–dark cycle). All monkeys were fed a daily diet of commercial marmoset food that was modified (Test Diet 5WW6) for high fiber and gum Arabic to support digestive health (Mazuri Marmoset Diet, St. Louis, MO), supplemented with fresh fruit and vegetables, nuts, mealworms, and caterpillars. Water was available ad libitum. All procedures accorded with the Guide for the Care and Use of Laboratory Animals and were approved by the NIMH Animal Care and Use Committee.

### Apparatus

All testing was conducted in a custom designed Plexiglass apparatus comprising three stimulus boxes (12“ x 12 x 12”) each connected to a central viewing area (30“ x 12” x 12”) assembled like a T-maze (Fig. 1). The stimulus boxes were adapted to connect to the animal’s home cage such that once securely mounted, a sliding door, when opened, allowed the monkey to enter the box and be safely transported to the testing apparatus. The stimulus box, containing the marmoset, was then attached to either end of the central viewing area, or to the middle as illustrated in Fig. 1-2. On the side opposite the sliding door was a wire mesh (8’’ x 8”) which restricted gross physical access but enabled social interaction by allowing animals to see, hear, and smell one another, as well as lightly touch each other through the holes in the mesh. A GoPro camera (Hero 9 Black; GoPro, San Mateo, CA) located above the apparatus provided a top-down view of the entire test compartment. The animals’ reactions to each other were scored manually.

### Behavioral Procedure

The animals were first habituated to the stimulus box by mounting it on the animals’ home cage. The monkey was allowed to freely enter and explore the box for 30 mins for three consecutive days. The females were then habituated to the love maze apparatus. The stimulus box containing the female was connected to the middle location of the apparatus; the stimulus boxes flanking each side of the central viewing area were empty. The female was then allowed to move around the enclosure for 30 mins for three consecutive days to acclimate to the new environment. On the day of testing, two stimulus boxes each containing a male, unfamiliar to the female, were connected to either end of the apparatus with the mesh facing the viewing area. Thus, each male could see each other across the maze but were never released into the viewing area. Following a 20-minute acclimation period for each male, the female test subject was brought to the apparatus in her stimulus box and attached to the middle location (**see Fig 1**). The sliding door was opened, and the female was allowed to enter the central viewing area and safely interact with the two males located behind the mesh at each end of the apparatus. Each test session lasted 30 mins.

Initially, each female started with two random unknown males to choose from. All sexual solicitations (e.g., proceptive tongue flicks, presentation, lip smacking, grooming) from either male or female were recorded. If the female showed no preference in either male, she was presented with a new pair of unknown males on the following day (**Fig. 2A**). The first session in which she showed a clear preference towards one male by showing sexual solicitations, that male went through two more sessions to confirm a consistent preference (**Fig 2A**). The left and right positions of the preferred male alternated between sessions. The female was required to show preference for the same male across three consecutive sessions. If she failed to show a consistent preference for one male across three sessions, additional males were added to the choice (**Fig. 2B**). Once the female showed a clear preference for one male in three consecutive sessions, a pair was established, and the two animals were paired in the same home cage for mating.

### Mating preference

Sociosexual, affiliative, aggressive/territorial, and proximity behaviors were observed and manually recorded for both males and females. Behavioral expressions of preference towards the male (e.g., time spent in close proximity, number of tongue flicks, lip smack, presentation, grooming) were compared between animals using descriptive statistics and analyzed for statistical significance using t-tests. Tongue flicks and lip smacking from the female directed to a male was used as the primary indicator for preference, and once exhibited, that male would move to the next session. Two independent raters observed the expressed behaviors during two separate test sessions to confirm that the behaviors exhibited by the monkeys could be readily identified and distinguished from each other.

To circumvent obstacles faced by conventional video tracking methods when applied to marmosets, we used deeplabcut (version 2.3.4) (40,41) to determine the position and facing of the female with respect to each male. We used a ResNet-50-based network (42,43) with default parameters for 200k training iterations. The test error of the final model was 3.35 pixels and the train error was 5.8 pixels when applying a confidence threshold of 90%. The preference of the female for proximity to the preferred male over the non-preferred male was analyzed by a paired t-test.

The proportion of females pregnant at 90 d was analyzed using a Chi-square test of independence and the trajectory of reproductive success over time by a Gehan-Breslow-Wilcoxon test.

### Ultrasonography to detect conception

To determine a successful mating pair, we tracked the number of days to conception via ultrasonography. Females were imaged weekly using ultrasonography (Vevo MD Imaging System, Fujifilm, Visualsonics) to detect first signs of pregnancy. Detection of pregnancy was determined by the transverse diameter of the uterus and uterus lumen. The uterus lumen opening could be detected on ultrasonography twelve to eighteen days after ovulation (44). After the first week of a visible uterus lumen opening, three more weeks of an increase transverse size were required to confirm viable pregnancy.

### Random pairs

Male-female couples that were randomly paired for breeding were extracted from available records in the NIMH Veterinary Medicine and Resources Branch. A total of 39 pairings over a period of four years were included for comparison. These pairings consisted of 36 females (2-9yrs) and 32 males (2-12yrs). Two females and six males were involved in more than one pairing.

## Acknowledgements

This research was supported by the Intramural Research Program of the National Institute of Mental Health (ZIA MH002951 and ZIC MH002952 to Y.C, and MH002924-15 to KA-W). We thank George Dold and Phuoc Pham from the NIMH Section on Instrumentation for customization and fabrication of the partner preference test for mate selection. We also thank Dr. Aaryn C. Mustoe from University of Nebraska Omaha for his expert advice on marmoset mating behaviors. We are indebted to the NIMH Veterinary Medicine and Resources Branch (VMRB) and Central Animal Facility (CAF) for their services in animal health, hygiene, and husbandry.

## CRediT

Conceptualization and methodology (YC, TMD), investigation (TMD, AJ), formal analysis (TMD, SPB), visualization (YC, TMD, SPB), resources (YC, KA-W), writing - original draft (YC, TD), Writing - review and editing (YC, TMD, SPB, KA-W).

